# Machine learning-based prediction of human structural variation and characterization of associated sequence determinants

**DOI:** 10.64898/2025.12.09.693295

**Authors:** Daven Lim, Runyang Nicolas Lou, Nilah Ioannidis, Peter H Sudmant

## Abstract

Structural variants (SVs) represent a major source of genetic diversity and play key roles in human disease and evolution. Yet, the extent to which local sequence context shapes the likelihood of structural variant formation remains poorly quantified. Here, we develop machine learning models to predict the occurrence of SVs across the human genome and characterize genomic determinants associated with their formation. We developed both a sequence only-based convolutional neural network (CNN) model as well as a random forest approach integrating diverse genomic annotations. Both models achieve high predictive performance individually (>90% AUROC) which can be further improved in an ensemble. The predictive ability of these models demonstrates that SV-prone regions can be accurately inferred from sequence context. Model interpretability techniques reveal key genomic contributors to SVs, including effects of sequence motifs such as microhomology and non-canonical DNA structures, as well as the presence of SV hotspots. We find that different classes of SVs exhibit distinct sequence determinants, with transposable elements and inversions displaying particularly unique signatures. Moreover, predicted SV probability correlates with allele frequency and gene functional constraint, indicating the potential utility of the model for variant effect prediction. These findings demonstrate that machine learning models trained on local sequence features can identify unstable genomic regions and provide a framework for quantifying SV susceptibility and SV variant effects in personalized genomics.

## INTRODUCTION

Structural variants (SVs) - including insertions, deletions, duplications, and inversions - are large-scale genomic rearrangements that represent a major source of genetic diversity in humans^1,2^. SVs can affect molecular and cellular processes which can cause genetic diseases, affect gene function, and contribute to evolution and trait complexity^3^. For example, de novo duplications at the 7q11.23 locus are strongly associated with autism spectrum disorder, and the reciprocal deletion causes Williams-Beuren syndrome^4^. Rare SVs can alter genes in neurodevelopmental pathways and contribute to schizophrenia^5^. SVs are also an important source of adaptive traits. For instance, increased copy number of salivary amylase genes (*AMY1*) is correlated with increased salivary amylase protein levels, and individuals from populations with high starch diets have been shown to have on average more *AMY1* copies ^6^ ^7^.

SVs arise through various sources, each leaving characteristic signatures in the local sequence context. A major source of SVs is non-allelic homologous recombination (NAHR), which occurs when highly similar low-copy repeats or segmental duplications misalign during meiosis, producing deletions, duplications, and inversions^8^. Transposable element activity, such as SINE/Alu and LINE/L1-mediated retrotransposition, also contributes substantially to human structural variation through insertion events. Furthermore, these sequences can serve as substrates for homology mediated rearrangements^9^. In addition, replication-associated processes such as fork stalling and template switching can generate complex duplications and rearrangements without requiring extended homology^10^. Many human SVs also originate from non-homologous end joining (NHEJ), a repair mechanism that rejoins double-strand breaks with minimal or no homology and frequently introduces short insertions or deletions at the junctions^11^. Breakpoint analyses have consistently shown enrichment of short microhomology tracts at SV junctions relative to random expectation, highlighting the role of microhomology-mediated repair and the importance of local sequence composition in predisposing genomic regions to structural instability^12^. Collectively, these mechanisms point toward sequence-dependent determinants of SV formation, motivating the use of computational models to quantify how local sequence features shape genome fragility.

Recent work in a eukaryotic pathogen and subsequently in *Arabidopsis thaliana* has shown that machine learning models can predict structural rearrangements from local genomic features (e.g. proximity to TEs and genes), demonstrating that eukaryotic genomes contain identifiable signatures of chromosomal instability^13^. However, these studies were limited to smaller and less complex genomes and focused on annotated genomic features instead of raw sequence. The degree to which such predictive signatures exist in the human genome and the extent to which local sequence context influences SV formation is also unclear. Furthermore, there are currently no quantitative frameworks to estimate the likelihood that SVs arise at specific genomic positions and the extent to which raw sequence content can serve as an accurate predictor.

Here, we develop machine learning models to assess SV susceptibility across a large, population-scale SV dataset generated from long-read haplotype-resolved genomes. We integrate an expanded genomic feature space that includes both raw sequence and diverse genomic annotations such as conservation metrics, base composition, neighboring genomic variation, repeat annotations, and gene-related features. By examining feature contributions and learned sequence determinants, we identify genomic and sequence factors associated with SVs. Predictive models of species-wide SV occurrence have the potential to support disease variant applications, advance our understanding of genome evolution, and guide future studies on the mechanisms of structural rearrangement.

## RESULTS

### Development of machine learning models to predict structural variants

Human structural variants are a diverse form of genetic variation spanning five orders of magnitude in length (**Fig 1A**). Nevertheless, SVs are known to be associated with distinct sequence features (e.g. stretches of homology). We set out to develop machine learning models capable of predicting the occurrence of SVs from DNA sequence and associated DNA features. We employed a population-scale SV dataset from Phase 3 of the Human Genome Structural Variation Consortium^14^ This is the first population-scale set of human genome assemblies resolved to near T2T completeness, encompassing 173,824 distinct variants discovered in 65 individuals (130 haplotypes) from diverse ancestries. We first set out to develop a sequence-only deep learning model for SV prediction utilizing a residual convolutional neural network architecture similar to models that have been developed to predict splice site usage^15^, gene expression^16,17^, mRNA expression^18^, and accessible genomic sites^19^. This approach estimates a probability that any given basepair location is the breakpoint of a structural variant given its adjacent flanking sequence (**Fig 1B, C**) (**Methods**). We also constructed a random forest-based SV prediction model in which the probability of an SV is determined from the relative distribution of sequence features from the nucleotide of interest within the context window. These features include the density of SNPs and neighboring SVs, GC content, constraint (PhyloP score), gene density, and mobile element density, among others (**Fig 1C, D**). Finally, we constructed an ensemble model in which the CNN predicted probability was incorporated as a feature into the random forest thereby combining both sequence content and genomic features into a single model (**Fig 1C**, see **Methods**). These methods were compared against a simple logistic regression approach which incorporated all of the features of the random forest. All models were trained and tested on 347,648 total samples (173,824 SVs and 173,824 background regions), with metrics reported on the test set which is not encountered during training (**Methods**).

**Figure 1.**
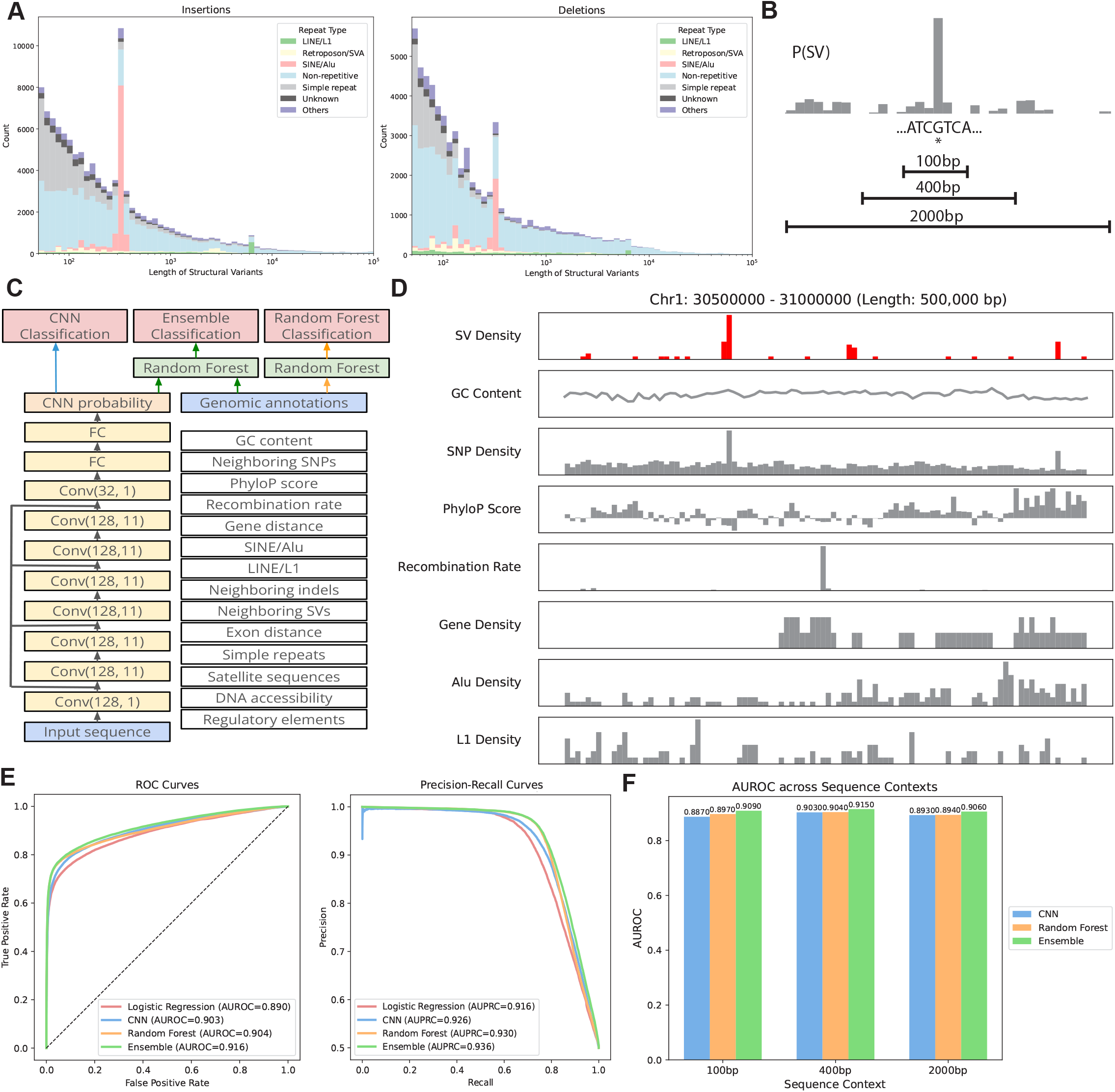
Details of the model and the prediction task. **(A)** Length distributions of insertions (left) and deletions (right) called against GRCh38 from HGSVC Phase 3, colored by repeat type. **(B)** Schematic of the model prediction task. At a target location in the human genome, models are trained to predict the probability of a structural variant event given sequence contexts of 100bp, 400bp, and 2000bp. **(C)** Schematic of the model architectures and predictions. **(D)** Genomic tracks along a 500kb region in chromosome 1 comparing SV density with a subset of genomic features. Bins of 5kb are used. **(E)** Receiver operating characteristic and precision-recall curves of the CNN, random forest, and the ensemble model, plotted against those of a baseline logistic regression model. **(F)** Comparisons of AUROC across sequence context lengths and model types.

We tested the performance of our models on the GRCh38 human genome equally split between true SVs and randomly selected control regions (**Methods**). Model performance was quantified using both the true positive and false positive rates assessed as the area under the receiver operating characteristic curve (AUROC) and the precision and recall rates, assessed as the area under the precision recall curve (AUPRC) (**Fig 1E, F**). All of our models outperformed the logistic regression approach, though this simple approach did perform surprisingly well, highlighting that local genomic features are strong determinants of the probability of SV formation (AUROC 0.890, AUPRC 0.916). The random forest approach modestly outperforms this basic method (AUROC 0.904, AUPRC 0.930), capturing non-linear relationships and higher-order interactions among genomic features that the linear logistic regression cannot represent. Remarkably however, the CNN approach, which only uses local sequence, achieved similar performance (AUROC 0.903, AUPRC 0.926). The ensemble approach combining the CNN with the random forest further improved performance (AUROC 0.916, AUPRC 0.936), and at a fixed 5% false positive rate, the ensemble exhibits a 6% improvement in true positive rate compared to the logistic regression, thereby quantifying the improvement in model performance through adopting a non-linear modelling approach and adding CNN predicted probability as a predictive feature. Sequence context lengths ranging from 100bp to 2000bp were also explored. The CNN and ensemble models were both improved by increasing the sequence context size up to 400bp, though diminishing beyond this point (**Fig 1F**).

To test our models on sequences outside the HGSVC3 dataset, we applied them to SV callsets from the Human Pangenome Reference Consortium (HPRC) that are generated from the same SV caller (PAV^20^) used by HGSVC3. To avoid contamination due to common variants present in both datasets, we held out SV calls from chromosomes 9, 10, 13, 17, and 21 (randomly chosen) and trained the models on HGSVC3 data without SV calls from these aforementioned chromosomes. We then evaluated model performance on 3 randomly chosen individuals from the HPRC dataset on SV calls from the held out chromosomes. The CNN achieves AUROC scores of above 0.86 and AUPRC scores of above 0.90, while the ensemble approach improves upon that, with AUROC scores of above 0.92 and AUPRC scores of above 0.94 (Fig S1). These results demonstrate the high performance of the models across varying data sets and contexts.

To further evaluate the performance of our models, we trained and tested them with the T2T-complete CHM13 reference genome using HGSVC3 SV calls for the CHM13 genome. As the CHM13 reference does not have the full set of genomic annotations, we tested only the CNN. On CHM13 with 188,500 SV (>50bp) calls, the CNN obtains an AUROC of 0.910 (GRCh38: 0.903) and AUPRC of 0.930 (GRCh38: 0.926), which is in fact slightly better than that obtained with the GRCh38 genome and could be attributed to improvements in assembly completeness and sequence accuracy.

Together, these results demonstrate that both genomic features as well as raw local genomic sequence are strong predictors of SV formation.

### Feature contributions to model predictions

Identifying and quantifying the impact of features to our model predictions provides insight into the mechanisms underlying SV formation. We conducted several analyses to quantify the contribution of individual features to model predictions. We first assessed the correlation between input features, the predicted SV probabilities, and SV presence/absence across models using Spearman correlation coefficients (**Fig 2A**). Several input features were strongly correlated with each other, as expected. For example, phyloP scores were negatively correlated with SNV and indel presence and DNase accessibility was correlated with transcription factor binding peaks. Feature correlations across models were highly consistent, with model probability strongly positively correlated with SNVs, indels, and the presence of other SVs, and negatively correlated with phyloP score and the distance to the nearest simple repeat. These features are similarly strongly correlated with the presence of SVs (“True label” **Fig 2A**). While it is expected that the random forest would pick up such features, surprisingly, the sequence-only CNN was also able to distinguish them. PhyloP score, for instance, exhibited a strong negative correlation with CNN output probability, indicating that the model is able to distinguish conserved regions of the genome from sequence alone.

**Figure 2.**
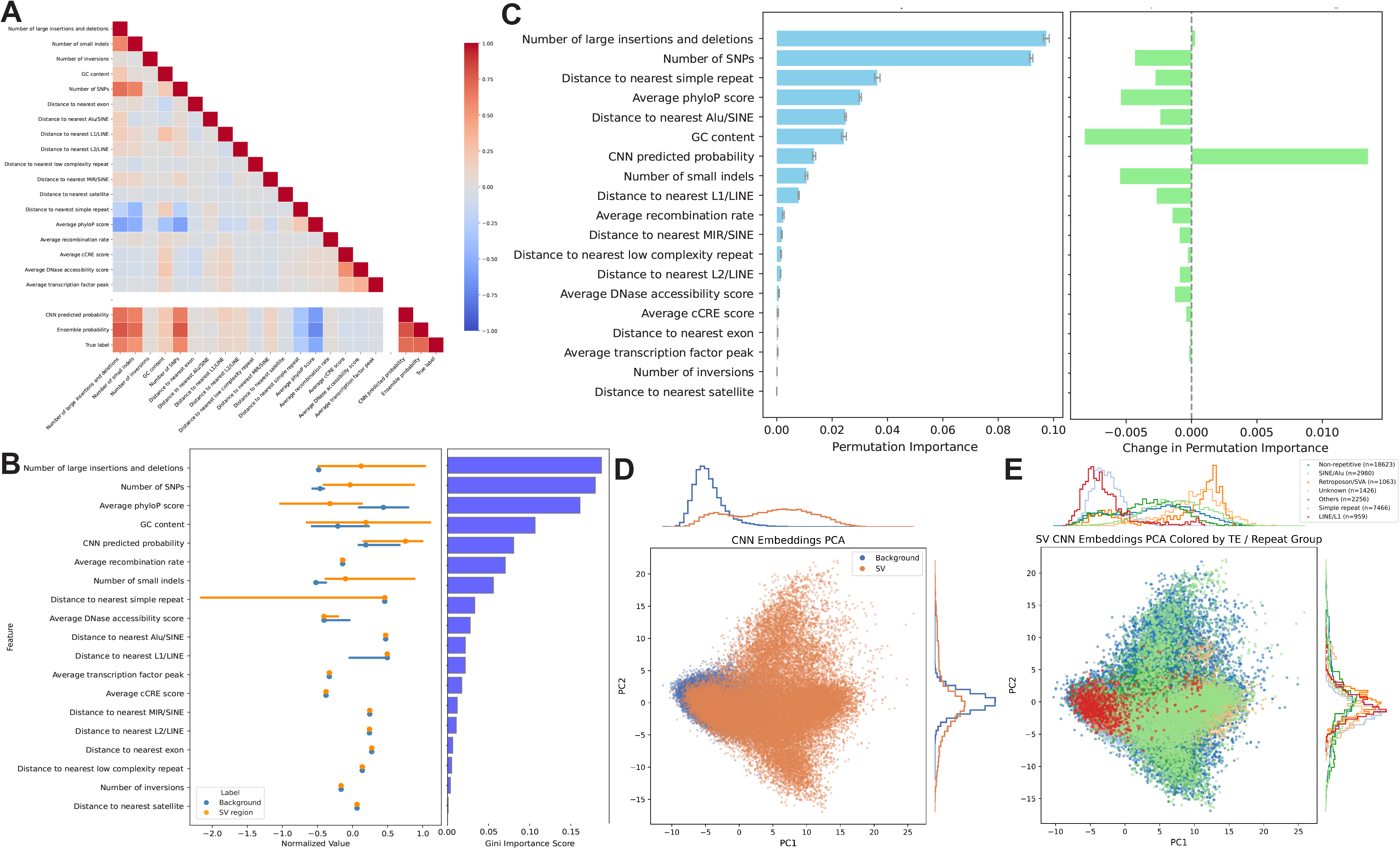
Analysis of model features. **(A)** Spearman correlation coefficients between genomic features, predicted SV probabilities, and true SV presence/absence of each model. **(B)** (Left) Distribution of genomic features used in the ensemble model in SV and background regions. Lines represent the interquartile range, and dots represent medians. Values are normalized to plot on the same scale. (Right) The Gini importances indicating the extent to which the model structurally relies on each feature for classification. Features are sorted with descending Gini importance from top to bottom. **(C)** (Left) Permutation importances of genomic features used in the ensemble model, which measures how much each feature affects model performance when randomly permuted. (Right) Change in permutation importance between before and after adding CNN predictions to the random forest model. **(D)** PCA of CNN final layer embeddings on the test set, colored by label. **(E)** PCA of CNN final layer embeddings on the test set, colored by SV subclass.

To determine the contributions of each feature to the model predictions, we first obtained the coefficients from the logistic regression (Fig S2). We find that neighboring variants - SVs, SNPs, and small indels - had the largest positive coefficients, suggesting that high density of neighboring variants is predictive of SVs, which aligns well with lower conservation scores serving as the next best predictor. Smaller distances to repetitive elements also rank highly in contributing to SV formation, most notably simple repeats.

We next evaluated the relative importance of each feature in the ensemble model, which includes the CNN-derived probability as an input (**Fig 2B**). Gini importance measures how much a feature reduces node impurity across all trees in the random forest, where impurity reflects how mixed the classes are within a node and a split that makes nodes more homogeneous decreases impurity. Distributions of each feature are shown alongside their Gini importances.

The number of neighboring SVs within the 400 bp window contributed most strongly to predictions, followed by the number of SNPs, conservation scores, and GC content. The CNN-derived prediction also ranked highly, indicating that sequence-level patterns captured by the CNN provide complementary predictive signals beyond genomic annotations.

To determine which labeled features the CNN is able to capture using raw sequence, we computed permutation importances in the ensemble model with and without the CNN as a feature on the test set. Permutation importances quantify the decrease in predictive performance when each feature is permuted. The decrease in predictive performance of a feature after permutation indicates the importance of the feature. Without the CNN as an input, neighboring variants - both SVs and SNPs - showed the highest contributions, followed by distance to simple repeats, conservation scores, GC content, and distance to Alu elements, largely consistent with relative feature importance measured with Gini impurity. This aligns closely with the feature coefficients in the logistic regression model. After introducing the CNN predictions as an additional feature, we observed a redistribution of importances, reflecting the extent to which the CNN captures signal previously explained by annotated features (**Fig 2C**). Conservation scores and GC content showed the largest reductions in importance, suggesting that the CNN effectively captures these sequence properties. The CNN also appeared to capture the number of SNPs and small indels, and the distance to simple repeats, Alu, and L1 elements. In contrast, the importance of neighboring large SVs increased, albeit slightly, indicating that local structural variant density provides information that is not fully captured by sequence context alone and remains a distinct predictor of SV likelihood. Although the CNN performs strongly on its own, its permutation importance is modest when added to the random forest, indicating that the ensemble is able to recover much of the same predictive signal from existing genomic annotations. The CNN therefore appears to provide more localized sequence-level cues in addition to the broader genomic features, such as short motifs and structural sequence patterns, which serves as a significant complementary signal.

We also performed further analyses on the CNN final layer embeddings on the test set. We plotted the embeddings in a 2D PCA space and observed that SVs occupy distinct spaces from background regions (**Fig 2D**). We also observed that SVs cluster within their subclasses (**Fig 2E**), discussed in a later section, and that SV sequences that overlap with background regions in the final layer embedding largely come from TE-associated sequences such as SINE/Alu and LINE/L1 (classification accuracy for SINE/Alu sequences: 0.503, for LINE/L1 sequences: 0.408), which is consistent with the small proportion of the total SV callset that TE-associated SVs form and that their insertion sites require the presence of short specific motifs instead of larger differences in sequence context. This also motivates our approach to subsequently model each SV subclass separately, to detect these specific sequence motifs without their signals being diluted by the larger number of non-TE associated SVs.

### Effect of microhomologies, non-canonical DNA structures, and motifs of varying composition

We next sought to identify the more fine-grained sequence features which the CNN identified as being associated with SVs. We performed a series of ablation analyses to quantify the effects of microhomologies, non-canonical DNA structures, and sequence motifs of varying composition on the predicted likelihood of SV formation. We first computed the enrichment of k-mers of specific lengths in SV regions relative to randomly sampled background sequences and identified k-mers that were significantly overrepresented near SV breakpoints. To test whether these enriched motifs influence model predictions, we inserted each k-mer at random positions within background 400 bp windows (**Methods**). For each modified sequence, we computed the change in predicted SV probability relative to the original unmodified background sequence and used the average change as a measure of the k-mer’s effect. We observed a positive correlation between k-mer enrichment (k=12) and the magnitude of prediction increase induced by insertion (**Fig 3A**), indicating that the CNN has learned sequence patterns associated with elevated SV susceptibility. To examine relationships among motifs with similar functional effects, we embedded k-mers (k=12) in a 2-dimensional representation space with DNABERT embeddings^21^ and found clusters of sequence-similar k-mers that produced comparable effects on predicted SV probabilities, reflecting the contributions of specific sequence patterns. We further identified a large cluster of k-mers that lead to a consistent positive net increase in predicted SV probability and examined the shared features of these k-mers. We found that they are significantly enriched in GC content and CpG islands, suggesting a shared mechanism leading to structural instability (**Fig 3B**).

**Figure 3.**
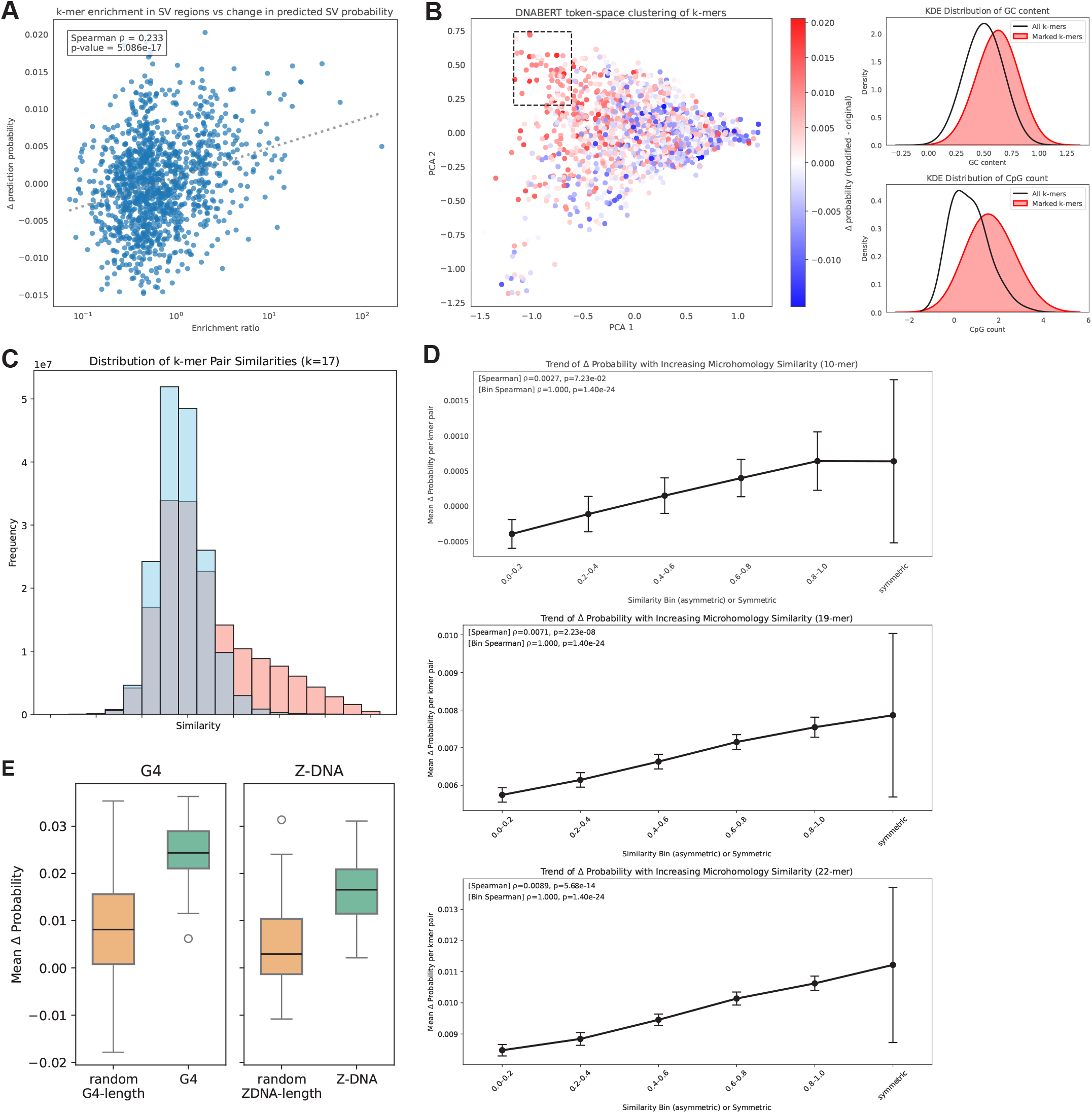
Effect of sequence motifs on CNN predictions. **(A)** Relationship between k-mer (length=12) enrichment in SV vs background regions, and the change in predicted SV probability post-insertion of each k-mer. A linear regression is shown as a dotted line, along with Spearman correlation and p-value. **(B)** (Left) DNABERT token-space clustering of k-mers of length 12, colored by change in SV probability post-insertion of each k-mer. The marked box shows a cluster of k-mers that all induce a positive change in SV probability. (Right) Density distributions of GC content and CpG count in the k-mers highlighted in the box and all plotted k-mers. **(C)** Empirical frequency of k-mer (length=17) pairs that share various degrees of similarity, across SV breakpoints (red) vs background locations (blue). Similarity scores increase from left to right. **(D)** Relationship between similarity between k-mer pairs inserted across a background location and mean change in predicted SV probability, for k-mers of length 10, 14, and 22. Similarities are binned, and fully symmetric insertions are included as the last bin. Spearman correlation and p-value are labelled as well. **(E)** Mean change in predicted SV probability after inserting G4-associated sequences vs. random sequences of G4-length, and Z-DNA-associated sequences vs. random sequences of Z-DNA-length.

Given the well-established role of microhomology in double-strand break repair and SV formation^12^, we next quantified the model’s sensitivity to short regions of breakpoint-spanning homology. Empirically, SV regions showed a higher frequency of k-mers with higher homology across breakpoints compared to background windows (**Fig 3C**). To determine if the CNN was able to detect such microhomologies, we then inserted pairs of k-mers symmetrically around the breakpoint at random distances from the center. These insertions changed the model probability regardless of the similarity between k-mers, indicating that the model is sensitive to artificial looking DNA sequences. Remarkably, however, increased homology between k-mers was significantly associated with an increase in predicted SV probability (**Fig 3D**). This effect increased in strength as a function of motif length. These results are consistent with the model capturing sequence features relevant to microhomology-mediated repair.

Finally, because non-canonical DNA structures have been observed to be enriched within the vicinity of SV breakpoints^22^, we assessed the model’s response to the insertion of sequence elements capable of forming G-quadruplex (G4) and Z-DNA structures. We inserted these motifs into background sequences, similar in approach to k-mer insertions, and compared the resulting change in predicted SV likelihood against random control sequences of matched length. Both G4 and Z-DNA insertions led to higher predicted SV probabilities (**Fig 3E**), suggesting that the CNN recognizes sequence features associated with structures that can increase fragility at SV breakpoints.

Distinct sequence signatures underlie different SV mechanisms, and together, these results highlight that the CNN model accurately captures these features associated with SVs. Furthermore, these features exhibit different scales; the CNN identified sequence interactions indicative of microhomology as well as individual motifs associated with SVs.

### Model performance across SV classes

SVs span a wide range of different sizes and are associated with many different mechanisms (**Fig S3**). We thus evaluated the performance of the models across different SV classes annotated with Repeatmasker: transposable element (TE) associated SVs, simple repeat associated, non-repeat associated, and inversions. For each subclass, we trained the CNN, random forest, logistic regression, and the ensemble model independently using the same procedures described above and compared their predictive performance.

Prediction accuracy varied markedly across SV types (**Fig 4A**). SVs associated with simple repeats were the most accurately predicted, with the CNN achieving an AUROC of 0.994, slightly outperforming the random forest (0.992), logistic regression (0.986), and the ensemble model (0.993). Similarly, TE-associated SVs showed strong sequence-driven predictability. For SINE/Alu SVs, the CNN reached an AUROC of 0.925, substantially higher than the random forest (0.614), logistic regression (0.626), and ensemble (0.807); LINE/L1 SVs showed a comparable pattern (CNN: 0.823; random forest: 0.676; logistic regression: 0.677; ensemble: 0.745). These results indicate that TE-mediated SVs are strongly characterized by local sequence signals rather than broader genomic context, and that sequence models can accurately predict TE-mediated SVs. In contrast, the CNN underperforms against the random forest model with non-repeat SV and in inversions. In particular, while none of the models performed particularly well on inversions, the CNN was extremely poor (0.480). These results highlight the varying importance of local sequence features to distinct SV classes.

**Figure 4.**
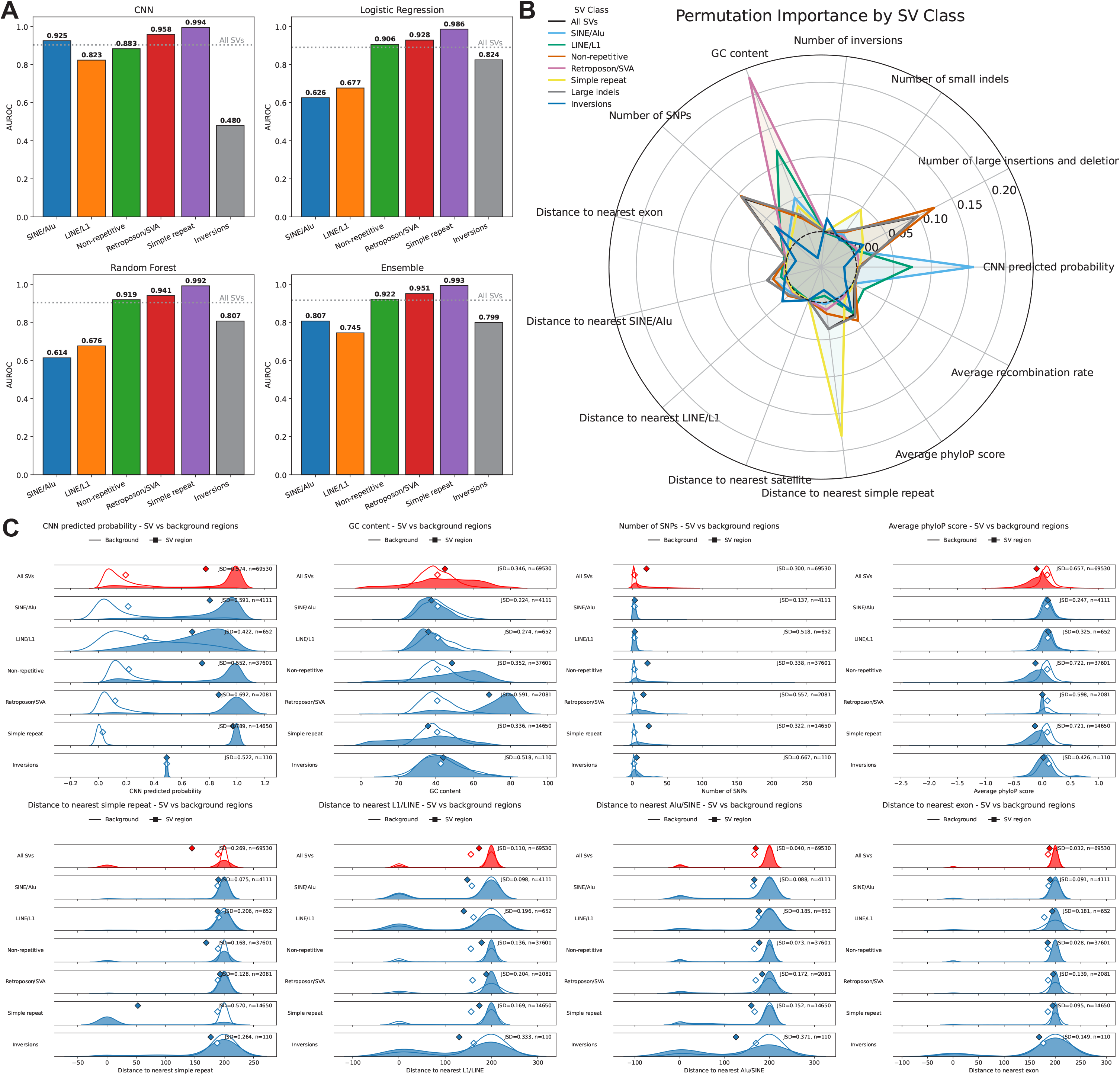
Model performance and feature distributions of SV subclasses. **(A)** Model AUROC performance across various subclasses and model types, with the dotted line representing baseline model performance for all SVs. **(B)** Permutation importance of features in the ensemble model for different SV subclasses. **(C)** Distributions of various features across SV and background regions, categorized by SV subclass. Feature distribution of all SVs are plotted in red for comparison. The diamond symbols represent the mean of the distribution.

To identify the most influential predictors for each SV class, we computed permutation importances for all features within each subclass and compared them to importance rankings for the full SV set (**Fig 4B**). CNN-derived sequence scores were among the strongest predictors for LINE/L1 and especially SINE/Alu associated SVs, reinforcing the role of TE-specific sequence patterns in driving these events. GC content was the dominant predictor for LINE/L1 SVs and was also highly informative for retroposon/SVA SVs, consistent with known compositional biases of these TE insertion targets. Distance to simple repeats was the strongest predictor for simple repeat-associated SVs, reflecting the tendency of these events to arise within repetitive regions^8^.

Inversions showed a notably different signature. Many features contributed with similar importance, and no single predictor dominated. The number of neighboring inversions emerged as a strong feature, suggesting the presence of inversion hotspots. Distance to LINE/L1 elements was also informative, consistent with mechanisms in which L1 retrotransposition facilitates rearrangements in adjacent sequences. The number of SNPs was additionally predictive. Interestingly, several features exhibited negative permutation importance values, suggesting that permuting these features slightly improved performance. This may indicate redundancy among highly correlated predictors or that certain features add noise rather than informative structure.

### Distinct sequence and genomic feature distributions across SV classes

To evaluate how sequence features differ between SV regions and background genomic windows, we compared the distributions of multiple genomic annotations across SV classes and computed the Jensen-Shannon divergence (JSD) between SV and background distributions (**Fig 4C, S4**). Higher JSD values indicate greater distributional separation and therefore stronger enrichment or depletion of that feature near SV breakpoints. Each feature includes the distribution of all SVs for comparison.

Across all SVs, breakpoint windows consistently harboured higher local variant density than background. SV regions contained more SNPs, small indels, and large insertions/deletions, with moderate-to-high JSD values for each (around 0.30-0.50), indicating that SVs tend to arise in already variant-rich regions, or that SV formation can induce local mutations, which has been previously observed^23^. This enrichment was particularly pronounced for inversions, which showed the largest divergence in SNP counts (JSD = 0.667). In contrast, the number of inversions in the surrounding window showed only modest shifts for most classes, but was clearly elevated around inversion breakpoints themselves (JSD = 0.196), supporting the presence of localized inversion hotspots.

Conservation and base composition also differed substantially between SV and background windows. Average phyloP scores were markedly lower at SV breakpoints than in background, especially for non-repetitive and simple-repeat SVs (JSD ≈ 0.7), consistent with reduced evolutionary constraint at rearrangement-prone loci. GC content likewise showed strong divergence for several classes, including retroposon/SVA events and inversions (JSD up to ∼0.6), indicating that GC-rich or GC-poor environments contribute differentially to specific SV mechanisms. This highlights that the importance of GC-content toward the likelihood of SV formation is dependent on SV class, and is most apparent for SVA elements which have been previously shown to insert in high GC and genic sequences^24^.

Overall, SVs occurred closer to simple repeats than background regions, with the largest effect for simple repeat-associated SVs themselves (JSD = 0.570), confirming that these events are highly concentrated within repeat-rich regions. Several class-specific biases can be observed - LINE/L1, SINE/Alu, and inversion SVs were found closer to LINE/L1 elements compared to background, and inversions were also enriched nearby SINE/Alu elements. By contrast, distances to the nearest exon showed relatively small JSD values across all classes. These comparisons reveal that different SV classes occupy distinct regions of genomic feature space. Overall, breakpoints are preferentially located in regions of elevated local variation, reduced conservation, characteristic GC content, and specific arrangements of repetitive elements, with particularly strong and class-specific signatures for repeat- and TE-associated SVs and for inversions.

### In silico saturation mutagenesis for TE insertions

The CNN model performed particularly well on TE insertions compared to other SV classes. To identify sequence features underlying the model’s performance, we performed in silico saturation mutagenesis (ISM) on a subset of held-out test sequences corresponding to Alu-, L1-, or SVA-associated SVs (**Methods**). This approach entails mutating bases within ±30 bp of the breakpoint of TE-associated SVs, one nucleotide at each position at a time. For each mutation, we then computed the change in the predicted CNN probability relative to the unmutated sequence. This process results in an averaged mutational sensitivity map representing sequence determinants of TE-associated SV susceptibility learned by the model (**Fig 5A, B**).

**Figure 5.**
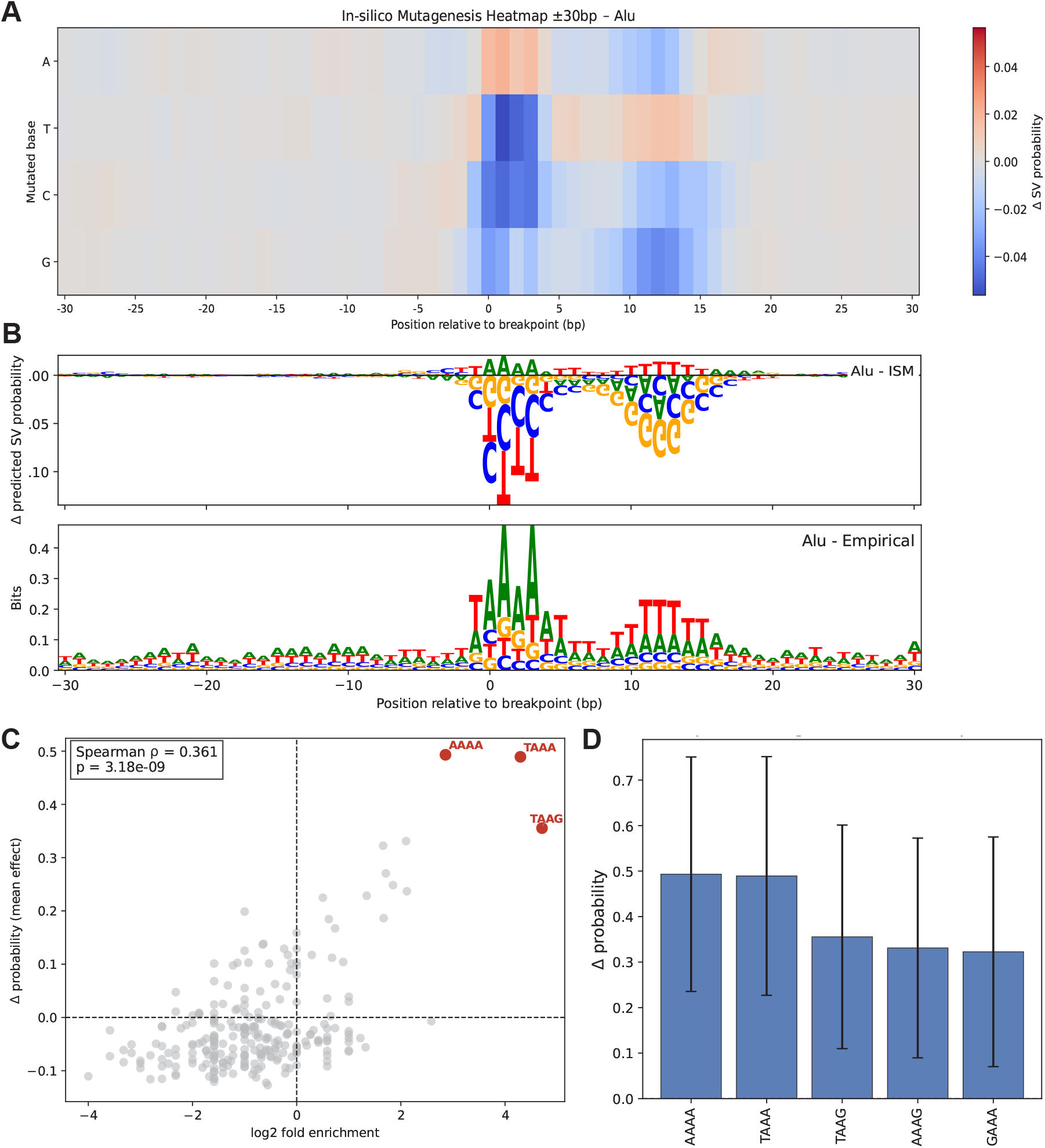
In-silico mutagenesis and empirical findings for Alu-associated SV sites in the test set. **(A)** ISM heatmap showing change in model predictions from mutating each nucleotide across ±30 bp, highlighting positions at which substitutions strongly modulate predicted SV likelihood. **(B)** (Top) ISM sensitivity depicted with sequence illustrations, where the height and direction of each letter reflect the magnitude and sign of the mutational effect. (Bottom) Empirical nucleotide composition computed directly from the unmutated test-set sequences, summarizing the observed frequency of each nucleotide at each position relative to the breakpoint. **(C)** Mean change in model predicted probability of inserting motifs at Alu-associated breakpoint sites against enrichment of motifs compared to background sequences. **(D)** Mean change in model predicted probability of inserting motifs at Alu-associated breakpoint sites, with error bars denoting standard deviations.

We find that the effect of ISM on TE-associated SV sites is highly localized, with model sensitivity concentrated within ∼15bp of the breakpoint. The model strongly favors TAAAA motifs starting at the −1 position relative to the breakpoint. This motif perfectly resembles the empirical sequence enrichment around breakpoints (**Fig 5B, Methods**). Previous work has detailed the TAAA L1-endonuclease cleavage motif that we observe^25^. We also observe a, subsequent T-rich stretch downstream of the breakpoint which likely results from asymmetry in the breakpoints identified by the SV caller around TE-associated polyA tails.

To extend beyond the impact of single base pairs we compared the change in the model probability after mutating 4bp motifs at the center of randomly selected background sequences against the enrichment of these motifs at Alu-associated TE breakpoints (**Fig 5C, D**) (**Methods**). These metrics are significantly positively correlated (ρ = 0.361, p=3.18e-09) indicating that the model probabilities are strongly reflective of sequence compositions that are commonly associated with Alu SVs. Indeed, the average change in probability for the top 5 motifs ranged from ∼0.3-0.5, highlighting the incredibly strong preference for insertions at the canonical target site. In addition, we ran DeepSHAP^26–28^ across a set of SVs from each class to assign importance scores to individual nucleotides or sequence features, and then TF-MoDISco to convert nucleotide-level SHAP scores into motifs to discover sequence patterns that the CNN associates with SV formation. As the positions of the motifs can matter significantly, particularly for TE-associated sequences, we further included a framework to look at motif scores by position to identify the positions of high-contribution motifs. To test the framework, we looked at closely-related motifs to the TAAA endonuclease cleavage site for Alu insertions across all SV subclasses, and find that TF-MoDISco^29^ on DeepSHAP attribution scores by position correctly identifies the TAAA motif at the correct position for Alu insertions, while not detecting similar signals across other SV subclasses (**Fig S5**). Together these results highlight the CNN model’s ability to capture highly specific and well-studied sequence features associated with SVs.

### Gene functional constraint and tolerance

Structural variants play key roles in human disease and phenotypic variation^3^ with strong selection observed against large SVs^1^. However, establishing the relationship between phenotype and genotype remains a key challenge in human genetics. Machine learning is providing opportunities for understanding these high-dimensional relationships^30^, but most existing methods focus on SNV^31^ while SVs are still underexplored. To assess the potential of machine learning models for predicting the functional impact of SVs, we explore how our model predictions interact with features such as allele frequency, genomic constraint, and variant effect predictions.

Deleterious SVs are subject to negative selection and are thus enriched for rare variants^1^. We thus examined the association between our model predictions and allele frequency. We note that the training and testing tests were randomly split, alleviating any allele frequency biases in the generation or testing of our model. Predicted SV probability increased with allele frequency with common SVs receiving substantially higher predicted probability than rare SVs (**Fig 6A**). This was true for both the CNN (p=1.6e-7) and ensemble models (p=1.2e-22), indicating that the model may be able to identify features related to SV persistence or genomic mutability in the population, thus favoring common SVs versus rarer ones. Indeed, phyloP score is strongly correlated to both CNN and random-forest predicted probabilities (**Fig 2A**).

**Figure 6.**
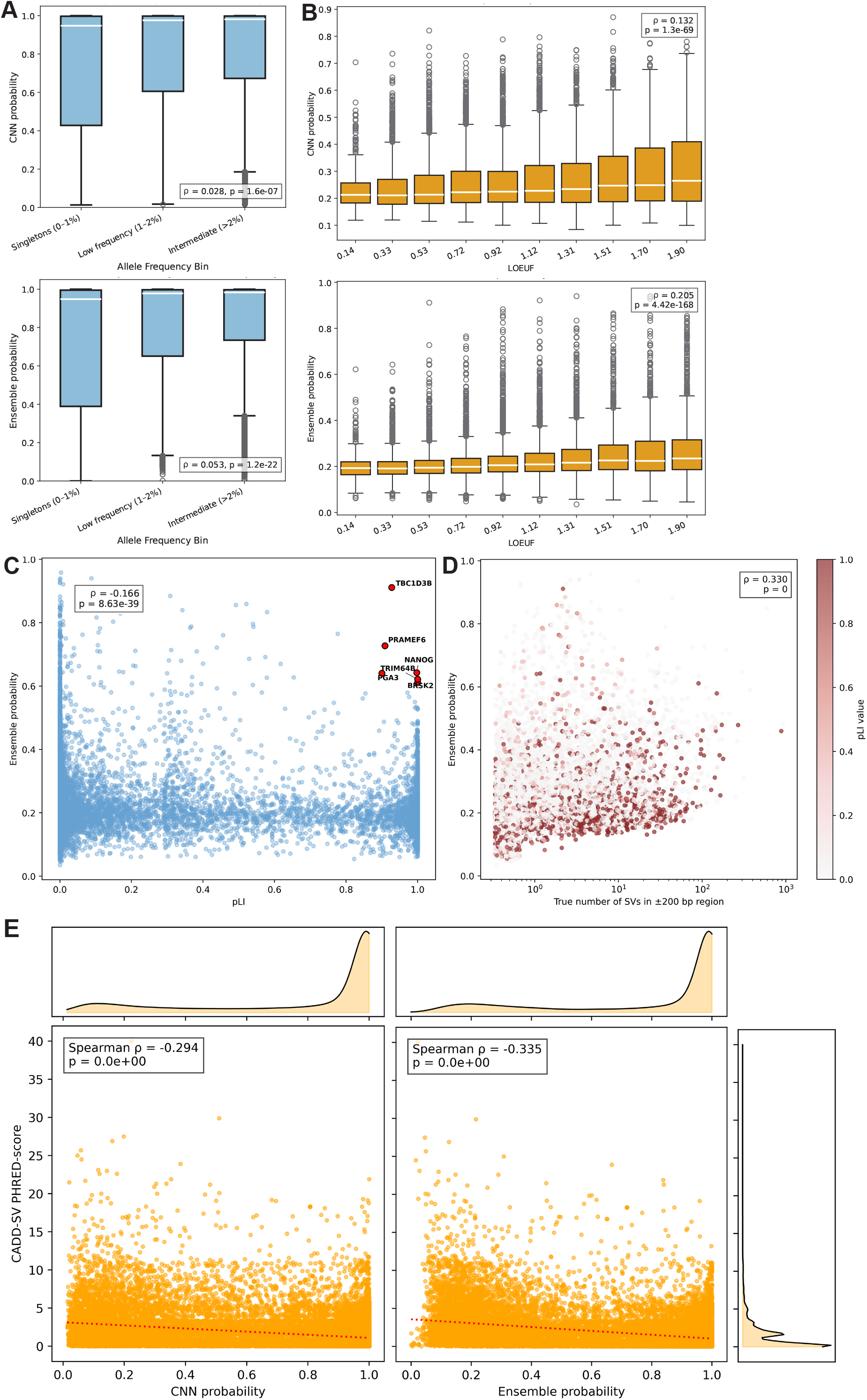
Interactions between model prediction and the selective constraint and functional impact of SVs. **(A)** Relationship between allele frequency in HGSVC3 and predicted SV probability of the CNN and ensemble model. Allele frequency is binned into singletons, low frequency variants, and intermediate variants. **(B)** Relationship between gene LOEUF and predicted SV probability of the CNN and ensemble. **(C)** Ensemble predicted probability plotted against pLI, with outlier genes that have high constraint and high predicted SV probability highlighted in red. **(D)** Relationship between ensemble predicted probability and observed SV count in each gene region, colored by pLI. Points are jittered slightly along the x-axis for visualization. **(E)** Correlation between CADD-SV PHRED-scores and model predictions from both the CNN and ensemble. Density plots are plotted for each axis. A linear regression is shown as a dotted line.

To further explore the relationship between constraint and our models, we examined model predictions in genes with constraint quantified using LOEUF (gene loss-of-function observed/expected upper bound fraction)^32^ and pLI (probability of loss intolerance)^33^. LOEUF increases as a function of relaxed constraint while pLI increases as a function of increased constraint. We sampled 30 random locations within each gene and calculated the averaged predicted SV probability. Model predictions over genes were strongly correlated with both LOEUF (**Fig 6B**, ρ = 0.205, p-value = 4.42e-168) and pLI (**Fig 6C**, ρ = −0.166, p-value = 8.63e-39) indicating the model is identifying features associated with SV persistence and likelihood. Indeed, model predictions over genes were correlated with the number of true observed SVs over those genes (**Fig 6D**, ρ = 0.330, p = 0). When genes were colored by their pLI values, we observed that even among constrained genes, some loci sit in SV-prone genomic neighborhoods. However, highly constrained genes tended to exhibit lower predicted SV probabilities. This suggests that beyond raw SV density, the model is able to distinguish intrinsic sequence fragility from the selective pressures that shape the persistence of SVs near essential genes.

Finally, we compared the relationship between CADD-SV variant prediction scores^34^ and our model predictions for all SVs in the held-out test set. CADD-SV scores reflect putative variant pathogenicity with higher scores indicating more deleterious impact. Both the CNN and ensemble models showed significant negative correlations with CADD-SV PHRED scores (**Fig 6E**, CNN: ρ = −0.294, p = 0; Ensemble: ρ = −0.335, p = 0) indicating that predicted deleterious SVs have lower probability in our model predictions. These observations show that the model has putative utility for variant effect prediction despite not being specifically designed for this purpose.

Our analyses did reveal several outlier genes and SVs showing markedly higher predicted SV probabilities than expected for their level of constraint or predicted variant impact (**Fig 6C, D, E**). These outliers indicate genomic regions where local sequence architecture is inherently more prone to structural instability despite strong functional constraint, suggesting possible decoupling between sequence fragility and selective pressure. An example is the *TBC1D3B* gene which exhibits extreme structural heterozygosity, with haplotypes differing by hundreds of kilobases and in gene copy number^35^. This gene plays important roles in human brain evolution and resides within highly variable segmental duplications. Similarly, Nanog genes are flanked by Alu elements that modulate their regulation and expression^36^. Nevertheless, such sequence features can predispose this region to SVs and highlight how certain loci may occupy genomic environments predisposed to recurrent SV formation despite being under substantial functional constraint.

## DISCUSSION

In this study, we demonstrate that SV susceptibility in the human genome can be predicted with high accuracy using local sequence context and genomic annotations. The strong performance of both the CNN and the random forest model, and the further improvement achieved by combining them in an ensemble, indicates that sequence-derived features contain substantial information about the likelihood of SV formation in human genomes. These results provide evidence that intrinsic genomic properties shape the distribution of SVs across the genome and demonstrate the potential to quantify this susceptibility at base-pair resolution.

Our feature interpretation analyses provide insight into the specific genomic properties associated with elevated SV likelihood. Correlation patterns were broadly consistent with established mechanisms of genome instability, where regions with lower evolutionary conservation and increased density of nearby genetic variants showed higher predicted SV probabilities, reflecting relaxed functional constraint and a greater tolerance for rearrangement. The strong influence of nearby SVs and SNPs, observed in both Gini and permutation importances, suggests that SVs tend to accumulate in genomic neighborhoods predisposed to breakage or rearrangement, forming local hotspot environments. This observation is supported by empirical evidence highlighting meiotic breaks as key drivers of SNV and SV formation^23^, as well as segmental duplications^1,37,38^. Importantly, our analyses of the CNN model demonstrate that local sequence patterns carry biological signals sufficient to identify such hotspots.

We performed extensive analyses to identify the features learned by the CNN model. These analyses indicated that CNN can effectively identify short sequence signatures that contribute to local fragility. These include microhomology sequences flanking SV breakpoints, as well as sequences capable of forming non-canonical DNA structures, such as G-quadruplex and Z-DNA, which can disrupt replication or repair processes. These findings suggest that the model captures the landscape of genomic fragility shaped by local sequence composition.

The likelihood of structural variation is also shaped by distinct and interpretable genomic signatures that differ across SV classes. The high predictive performance for repeat-associated SVs indicates that these events are strongly driven by local sequence composition and recognizable motif patterns, rather than broader genomic context. This reinforces mechanistic models in which polymerase slippage and retrotransposition generate rearrangements at sequence-encoded instability sites. In contrast, inversions displayed much lower CNN performance, suggesting that their formation depends less on short motifs and more on larger-scale genomic architecture. For TEs specifically, we find that the CNN model is able to capture known sequence features. Specifically, the CNN identified the well-characterized L1-endonuclease cleavage motif and additional associated sequence. In contrast, the random forest-based approach exhibited substantially less power to identify Alu and L1 insertions.

These findings suggest that the overall human SV landscape emerges in part from two interacting forces - locally encoded fragility (driven by motifs, base composition, and repeats), and regional tolerance for rearrangement (reflected in variant-rich and weakly conserved contexts). This framework explains why sequence-based models are accurate at predicting repeat- and TE-driven SVs but perform less effectively for inversions, which depend on long-range homology not captured within a 400 bp window. By demonstrating class-specific predictability and genomic determinism, we provide a foundation for SV risk annotation and suggest that integrating broader structural context may further improve predictions for inversion and other non-local SV mechanisms.

The relationship between model predictions, SV allele frequency, and gene constraint provides insight into how the human genome is organized into regions of structural stability and fragility. The higher predicted probabilities assigned to common SVs suggest that these variants arise in genomic environments that are both intrinsically prone to rearrangement and evolutionarily permissive to their persistence and that common SVs mark fragile sequence contexts. Regions near LoF-tolerant genes showed higher predicted SV probabilities than those near constrained genes. This association implies that the local sequence environments surrounding functionally less essential genes tend to harbor the structural and compositional features linked to SV formation. These findings suggest that the genomic landscape reflects a degree of coupling between functional importance and structural permissiveness. However, this association, though highly significant, is weak.

The overall predictive framework introduced here has several implications. First, it provides a quantitative tool to estimate SV susceptibility at scale and identify functionally constrained genes that are in SV-prone regions. Thus, these models can support studies seeking to understand putatively pathogenic rearrangements and in the construction of SV mutability maps of the human genome. Such mutability maps can be constructed for personalized genomes, as well as on datasets of disease-causing variants to determine if they occur in highly mutable loci.

Next, the identification of specific sequence determinants can inform mechanistic studies into SV formation pathways. In particular, our models can be employed on data from other species to understand better how genome structure evolves. In addition, the relationship between predicted SV probability, observed SV allele frequency, and functional constraints suggests that the model highlights regions where rearrangements not only arise but also persist within human populations, pointing to a potential role in long-term genome evolution.

Our study is not the first to employ machine learning approaches towards the prediction of SVs from genomic data^13^. However, our approach both improves upon previous work and exhibits several novel elements. For instance, we comprehensively compare several distinct predictive models including a CNN which only uses sequence data. We also perform prediction SV subclasses analyzing transposable elements, repeats, and inversions separately to characterize the differences in model predictive performance among these classes. Our study further examines a much larger range of SV sizes compared to previous work^13^. Importantly, large SVs are more likely to have high functional impacts and medical relevance. We also utilize finer-grained encoding of features. Instead of simple presence/absence encoding, we encoded counts, distances, rates, and scores of various features into our model resulting in more informative and interpretable results. Together, the many improvements in our models, alongside our use of human data, make them substantially more useful for personal genomics and translational applications.

Future work may extend this approach to additional structural variant types and to long-read sequencing data that capture a broader spectrum of rearrangements. Integrating replication timing, chromatin organization, or 3D genome architecture may further improve predictions, particularly for complex events. Additionally, a major limitation of our models is that they incorporate only highly localized sequence features, contributing to their failure to predict specific classes of SVs accurately (e.g. inversions). Accounting for both long and short-range sequence features and homology such as the usage of attention-based models has the potential to vastly extend the utility of our approaches. Finally, beyond the human genome, applying our framework across species may reveal species-specific and conserved determinants of chromosomal instability providing insights into the evolutionary forces shaping genome structure.

Overall, this work provides a genome-wide framework for quantifying SV susceptibility at nucleotide resolution. The models and analyses presented advance our understanding of the sequence architectures that predispose the genome to rearrangement, offering potential applications in variant interpretation, disease gene discovery, and the identification of rare but high-risk loci relevant to personalized genomics.

## METHODS

### Data sources

Structural variant calls were obtained from Phase 3 of the Human Genome Structural Variation Consortium (HGSVC3), using variants reported against the GRCh38 reference genome. Variant callsets included large insertions and deletions, large inversions, small indels, and SNPs. In the various callsets, there are 111,803 large insertions, 64,428 large deletions, 300 inversions, 3,128,354 small insertions, 3,049,659 small deletions, and 23,462,179 SNPs.

Genomic annotations were retrieved from the UCSC Genome Browser, including phyloP conservation scores, recombination rate estimates, RepeatMasker repeat annotations, candidate cis-regulatory elements (cCREs), DNase accessibility, and transcription factor ChIP– seq peak clusters. Reference genome sequences and gene annotations were obtained for GRCh38.

Further benchmarks for the model were made using variants reported against the CHM13 reference which can also be found in HGSVC3, and variants in 3 individual genomes (HG00658, HG01099, HG02027) from the Human Pangenome Reference Consortium (HPRC).

HGSVC callsets (for both GRCh38 and CHM13): https://ftp.1000genomes.ebi.ac.uk/vol1/ftp/data_collections//HGSVC3/release/Variant_Calls/1.0/GRCh38/

HPRC callsets: https://s3-us-west-2.amazonaws.com/human-pangenomics/index.html?prefix=submissions/759B21AD-0ED84640-A433-7C92A57EA3D3--UW_EEE_SV_Calls/

### Dataset construction

The final dataset consisted of 173,824 SVs, paired with an equal number of randomly sampled background genomic locations. Only fully contiguous 400bp sequences are included in the dataset. For each SV, we extracted a 400 bp context window centered on the breakpoint, comprising 200 bp upstream and 200 bp downstream (**Fig 1B**). For deletions and inversions, the internal deleted sequence was omitted from the window, where only the flanking sequences were used.

Overall, this yielded 347,648 total samples (173,824 SVs and 173,824 background regions). The dataset was partitioned into 60% training (n=208,588), 20% validation (n=69,530), and 20% test sets (n=69,530), with no genomic positions shared across splits. All model selection and hyperparameter optimization were performed on the training and validation sets. Only the test set was used for final evaluation and was never seen during training. Genomic annotation features - including conservation, GC content, repeat proximity, local variant density, gene features, and regulatory annotations - were calculated or extracted for each window. Sequence windows were one-hot encoded over four channels (ATCG). Spearman correlations across all features, model predictions, and true labels were calculated with SciPy.

To ensure that our approach does not introduce biases for deletions and inversions, we ran an additional analysis to sample breakpoints reflecting their true occurrences. Background windows for insertions consisted of the same 400 bp flanks surrounding the control positions. To construct inversion-specific negative examples, we generated simulated inversions matched in genomic length to the real events. First, we derived an empirical inversion length distribution from the set of true inversions and randomly sampled an inversion length for each negative example. A chromosome was selected at random, after which we sampled a first breakpoint uniformly along the chromosome followed by a second breakpoint, ensuring that neither the first breakpoint nor the corresponding second breakpoint (computed as first breakpoint + sampled inversion length) overlapped any known SV breakpoint. For each simulated inversion, we extracted a single 400 bp sequence window consisting of 200 bp immediately upstream of the first breakpoint and 200 bp immediately downstream of the second breakpoint, mirroring the windowing strategy used for true inversions. The internal sequence between the two breakpoints was excluded to match the representation of real inversion events. Genomic annotation features were computed over the two flanking windows and combined, producing negative examples that preserve the chromosomal context, breakpoint structure, and inversion length distribution of true inversions while ensuring no overlap with annotated SVs. Background windows for deletions were generated with the same method. We find that with this alternative approach for generating background sequences, there is little difference in the results we obtained.

### Convolutional neural network architecture

Sequence classification was performed using a custom 1D residual convolutional neural network (CNN) implemented in PyTorch 2.0.1. The model accepts a 4-channel one-hot– encoded 400 bp sequence. The architecture consists of an initial convolutional block of 1×1 convolution projecting 4 channels to 128, followed by batch normalization and ReLU activation. This is followed by a stack of 3 residual blocks, each consisting of Conv1d (kernel size 11, padding 5), BatchNorm1d, ReLU, second Conv1d (kernel size 11, padding 5), residual skip connection, and final ReLU. A 1×1 convolution then reduces dimensionality to 32 channels. The flattened output (12,800 units) is passed through a linear layer (128 units, ReLU), followed by a final linear layer mapping to a single output neuron. A sigmoid activation produces a probability estimate of SV occurrence.

The network was trained with binary cross-entropy loss, AdamW optimizer (with learning rate 5×10⁻ ⁶, weight decay 1×10⁻ ³), and a ReduceLROnPlateau scheduler (with factor 0.5, patience 3). Training used early stopping with patience of 5 epochs. The model was trained with batch-based stochastic optimization of batch size 128. The weights corresponding to the lowest validation loss were selected for all analyses.

### Random forest, logistic regression, and ensemble models

In addition to the CNN, we trained a random forest classifier using scikit-learn 1.6.1, a logistic regression baseline, and an ensemble model, implemented as a random forest receiving both genomic annotation features and the CNN-predicted probabilities as input to improve predictive performance.

For conventional machine-learning models, we combined the training and validation sets for model fitting after CNN hyperparameter selection. All models were evaluated on the held-out test set. Random forests used 1000 trees, balanced class weights, maximum number of features when splitting as the square root of the total number of features, and default bootstrapping. Logistic Regression was trained with L2 regularization.

### Feature importance analysis

Two complementary measures of feature importance were used - Gini-based importance, obtained from ‘sklearn.ensemble.RandomForestClassifier.feature_importances_’, and permutation importance, computed using ‘sklearn.inspection.permutation_importance’, which measures the decrease in model performance when individual features are permuted. Permutation importance was calculated on the test set to provide an out-of-sample estimate, with 5 repetitions. For analyses involving the ensemble model, the CNN probability was treated as an additional feature and its importance was evaluated alongside annotation features.

### K-mer, motif, microhomology, and non-canonical DNA structure analyses

To quantify the contribution of specific sequence patterns to SV likelihood, we performed targeted insertion experiments using the trained CNN model. All analyses were conducted on background (non-SV) 400 bp windows to ensure that observed prediction changes reflected the inserted sequence element rather than surrounding SV-associated context.

K-mer insertion analysis: We evaluated the effect of short sequence motifs by inserting all 1,024 permutations of 5-bp k-mers into 50 random positions within each background window. After each insertion, the sequence was truncated at the ends to maintain the fixed 400 bp window centered on the reference position. For each k-mer, we computed the average change in predicted SV probability across all insertions as a measure of its influence on model predictions.

Microhomology analysis: To assess the effect of short homologous tracts, we inserted 100 pairs of k-mers - either perfectly matching (symmetric microhomology) or mismatched (asymmetric controls) - symmetrically around a background location at 50 random distances from the center. As with the k-mer analysis, sequence windows were truncated to 400 bp after insertion. The change in predicted SV probability was averaged across all inserted microhomology pairs to quantify the model’s sensitivity to breakpoint-spanning homology.

Non-canonical DNA structure insertion: We next tested whether sequence elements capable of forming non-B DNA structures influence predicted SV likelihood. We inserted 100 distinct sequences predicted to form G-quadruplexes (G4) and 100 predicted Z-DNA -forming sequences into background windows at 50 random positions each. To ensure proper controls, we also generated 100 random sequences matched in length each to the G4 and Z-DNA motifs and inserted these in parallel. All modified sequences were truncated to retain a fixed 400 bp window centered at the reference site.

For all analyses, the mean change in CNN-predicted probability relative to the unmodified background sequence was used as the quantitative measure of motif effect size.

### Model construction for SV subclasses

Repeat annotations are performed using RepeatMasker version 4.1.5 on the SV callset. We label the SV with the repeat annotation if the repeat covers at least 80% of the SV sequence. In the case of multiple repeat annotations per SV, the repeat that spans at least 80% of the repetitive content in the SV sequence is utilized. Otherwise, the SV is labelled non-repetitive.

To evaluate whether different structural variant classes exhibit distinct sequence determinants, we trained separate models for individual SV subclasses. SVs were grouped into six categories: inversions, SINE/Alu-associated SVs (n = 4,111), LINE/L1-associated SVs (n = 652), non-repetitive SVs (n = 37,601), Retroposon/SVA SVs (n = 2,081), simple-repeat SVs (n = 14,650), and inversions (n = 300). For each subclass, we constructed a balanced dataset in which the positive set consisted exclusively of SVs from the focal subclass, and the negative set consisted of an equal number of randomly sampled background locations drawn from regions accessible in the reference genome. Context windows and annotations were extracted following the same procedures described for the full dataset.

For each subclass, we trained the same set of models used in the main analysis - the CNN, the random forest, logistic regression, and the ensemble model integrating the CNN prediction as an additional feature. The training, validation, and test splits followed the same strategy as for the full dataset, with no shared samples across splits. Model performance was assessed using AUROC and additional standard metrics. For interpretability, we computed permutation importances on the ensemble model with the test set for each SV subclass to identify the most influential genomic features and to characterize class-specific sequence determinants.

### In silico saturation mutagenesis for TE insertions

For each TE-associated SV in the test set, we extracted a 400 bp window centered on the annotated breakpoint. These windows represent the pre-insertion genomic context, and filtering by specific TEs isolates target-site features. For Alu insertions, we filtered for only full-length insertions using the most populated statistical histogram bin (n=1367). At each of these positions, we generated four mutated sequences by substituting the reference base with each alternative nucleotide (A, C, G, T), leaving all other positions unchanged. All sequences were one-hot encoded before passing through the trained CNN for each TE class.

To compute the effect of specific motifs at breakpoints of TE insertions, we inserted in-place 4-mers in the middle of background sequences and calculated the difference in model predictions pre- and post-modification. Specifically, we inserted the 4-mer to the left of the breakpoint to position the second nucleotide of the 4-mer at the breakpoint site, to mirror the effect of the T/AAA motif empirically observed at Alu-insertion sites. We then took the average of the effects across all sequences.

To calculate the empirical sequence enrichment around breakpoints, the process begins by separating sequences into positive sequences (e.g. Alu-associated breakpoints) and negative sequences (background/non-breakpoint regions). Each sequence has the same fixed length and is centered on the candidate breakpoint. For every position relative to the breakpoint, we count how often A/T/C/G occurs across all positive sequences, convert these counts into frequencies, and produce an empirical nucleotide distribution. This is repeated independently for the background sequences, generating a background nucleotide frequency distribution at every relative position. The enrichment calculation divides the positive frequency distribution with that of the negative, and measures whether a nucleotide is overrepresented near breakpoints compared to expectation from background genomic sequence.

### Constraint and tolerance

Gene constraint analyses used the gnomAD v4.0 gene constraint dataset, specifically the updated loss-of-function observed/expected upper bound fraction (LOEUF) metric and the probability of being loss-of-function (LoF) intolerant (pLI) score. Genes were matched using gene names from the gene annotation file and the gnomAD dataset. LOEUF and pLI values were matched to quantify how local constraint relates to predicted SV probability. Spearman correlations were calculated with SciPy. We quantified the number of SVs within each gene’s surrounding region (±200 bp).

### Software and reproducibility

All analyses were conducted in Python 3.10.14, random seed of 42, PyTorch 2.0.1, scikit-learn 1.6.1, transformers 4.41.2, SciPy 1.14.1, RepeatMasker version 4.1.5, and standard scientific libraries (NumPy, pandas, matplotlib). All code and processing pipelines were executed on GPU-enabled hardware when available.

## ACKNOWLEDGEMENTS

This work was supported by NIH National Institute of General Medicine award R35GM142916 to PHS. This work was also supported by the UC Berkeley Summer Undergraduate Research Fellowship (SURF) to DL. This manuscript is the result of funding in whole or in part by the National Institutes of Health (NIH). It is subject to the NIH Public Access Policy. Through acceptance of this federal funding, NIH has been given a right to make this manuscript publicly available in PubMed Central upon the Official Date of Publication, as defined by NIH. The content is solely the responsibility of the authors and does not necessarily represent the official views of the National Institutes of Health.

## AUTHOR CONTRIBUTIONS

PHS, NI, and DL conceived of the study. DL performed all analyses. DL, RNL, and PHS wrote the manuscript.

## DATA AND CODE AVAILABILITY

https://github.com/sudmantlab/sv_prediction.git

